# A chromosome-level assembly of the black tiger shrimp (*Penaeus monodon*) genome facilitates the identification of novel growth-associated genes

**DOI:** 10.1101/2020.05.14.096073

**Authors:** Tanaporn Uengwetwanit, Wirulda Pootakham, Intawat Nookaew, Chutima Sonthirod, Pacharaporn Angthong, Kanchana Sittikankaew, Wanilada Rungrassamee, Sopacha Arayamethakorn, Thidathip Wongsurawat, Piroon Jenjaroenpun, Duangjai Sangsrakru, Rungnapa Leelatanawit, Jutatip Khudet, Jasper J. Koehorst, Peter J. Schaap, Vitor Martins dos Santos, Frédéric Tangy, Nitsara Karoonuthaisiri

## Abstract

The black tiger shrimp (*Penaeus monodon*) is one of the most prominent farmed crustacean species with an average annual global production of 0.5 million tons in the last decade. To ensure sustainable and profitable production through genetic selective breeding programs, several research groups have attempted to generate a reference genome using short-read sequencing technology. However, the currently available assemblies lack the contiguity and completeness required for accurate genome annotation due to the highly repetitive nature of the genome and technical difficulty in extracting high-quality, high-molecular weight DNA in this species. Here, we report the first chromosome-level whole-genome assembly of *P. monodon*. The combination of long-read Pacific Biosciences (PacBio) and long-range Chicago and Hi-C technologies enabled a successful assembly of this first high-quality genome sequence. The final assembly covered 2.39 Gb (92.3% of the estimated genome size) and contained 44 pseudomolecules, corresponding to the haploid chromosome number. Repetitive elements occupied a substantial portion of the assembly (62.5%), highest of the figures reported among crustacean species. The availability of this high-quality genome assembly enabled the identification of novel genes associated with rapid growth in the black tiger shrimp through the comparison of hepatopancreas transcriptome of slow-growing and fast-growing shrimps. The results highlighted several gene groups involved in nutrient metabolism pathways and revealed 67 newly identified growth-associated genes. Our high-quality genome assembly provides an invaluable resource for accelerating the development of improved shrimp strain in breeding programs and future studies on gene regulations and comparative genomics.

## Introduction

Aquaculture is an important food source for the world’s growing population as it relieves the over-consumption pressure of natural animal populations within the aquatic environment. Currently, aquaculture provides the planet with more than 50 percent of fish products consumed by humans, making it one of the world’s fastest-growing food sectors with an annual growth rate of 5.8 percent since 2010 (FAO, 2018). The penaeid marine shrimps (Family Penaeidae) are the predominately cultured group (Thornber et al., 2019), with an annual production exceeding 4.5 million tons (Anderson, 2019). In this group, the Pacific white shrimp (*Litopenaeus vannamei*) and black tiger shrimp (*Penaeus monodon*) are the most dominant species cultured, accounting for 53% and 9% of total crustacean production, respectively (FAO, 2018).

While the penaeid industry has seen dramatic growth for the past few decades, industrial production of *P. monodon* proved to be unsustainable due to a lack of biological and genetic knowledge to achieve its desirable traits such as fast growth, disease resistance, reproductive maturation without reliance on wild brooders (Guppy et al., 2018). Recently, the *L. vannamei* breeding and domestication programs can be expeditely improved due to the available genome sequence, which allows selective breeding and helps in overcoming industrial challenges (Zhang et al., 2019). Genomic-driven breeding and domestication programs for *P. monodon,* on the other hand, are still in their infancy due to the absense of an informative high-quality draft genome sequence.

While such a high-quality draft genome sequence has not been reported for *P. monodon*, several efforts have been made to investigate the genome and genetic architecture of this important species over the past few decades. BAC library construction (Wuthisuthimethavee, Aoki, Hirono, & Tassanakajon, 2009), fosmid library end sequencing (Huang et al., 2011), molecular marker development (Brooker, Benzie, Blair, & Versini, 2000; A. Tassanakajon, Pongsomboon, Jarayabhand, Klinbunga, & Boonsaeng, 1998; A. Tassanakajon, Pongsomboon, Rimphanitchayakit, Jarayabhand, & Boonsaeng, 1997), linkage map construction (Baranski et al., 2014; Wilson et al., 2002), and transcriptomic analysis (Karoonuthaisiri et al., 2009; Leelatanawit, Uawisetwathana, Klinbunga, & Karoonuthaisiri, 2011; Lehnert, Wilson, Byrne, & Moore, 1999; Pootakham, Uengwetwanit, Sonthirod, Sittikankaew, & Karoonuthaisiri, 2020; Sittikankaew et al., 2020; Supungul et al., 2002; Anchalee Tassanakajon et al., 2006; Tong, Lehnert, Byrne, Kwan, & Chu, 2002; Uengwetwanit et al., 2018; Wongsurawat et al., 2010) were explored with limited success. Previous attempts to obtain genome sequences in the black tiger shrimp relied primarily on short-read sequencing platforms due to cost and technical difficulty in extracting high-quality, high-molecular weight DNA in this species (Quyen et al., 2020; Yuan et al., 2018). Recently, two draft genomes for *P. monodon* were made available; however, both versions of the assemblies were highly fragmented, with N50 contig lengths of merely 937 nt (Yuan et al., 2018) and 1,982 nt (Quyen et al., 2020). Even though those resources were useful for the black tiger shrimp genetics, they lacked the contiguity and completeness required for accurate genome annotation and thorough comparative genomics analyses.

Here, we combined a long-read sequencing technology and two long-range scaffolding techniques to obtain high-quality, chromosome-scale genome assembly. First, the Pacific Biosciences (PacBio) sequencing platform was employed to generate the preliminary assembly. The PacBio sequencing technology enables contiguous assembly of repetitive regions containing transposable elements and tandem repeats, which are often omitted or highly fragmented in genomic sequences currently available in public databases. We subsequently applied the long-range Chicago (*in vitro* proximity ligation) and Hi-C (*in vivo* fixation of chromosomes) scaffolding techniques to further scaffold the preliminary assembly to achieve the first chromosome-scale genome assembly in *P. monodon*. The utility of this dramatically improved assembly in elucidating novel genes involved in growth regulation was demonstrated by the transcriptome analysis of slow-growing and fast-growing black tiger shrimps. Our high-quality, chromosome-scale genome assembly provides a valuable genetic resource for black tiger shrimp breeding programs and future gene expression and comparative genomics studies in this species.

## Materials and Methods

### Sample collection and DNA extraction

The muscle of a 5-month-old female *P. monodon* was collected from the Shrimp Genetic Improvement Center (SGIC, Surat Thani, Thailand), immediately frozen in liquid nitrogen and stored at −80 °C until use. Frozen muscle was pulverized in liquid nitrogen and genomic DNA was extracted using a Genomic Tip 100/G kit (Qiagen, USA). DNA quantity was measured using a NanoDrop ND-8000 spectrophotometer and a Qubit dsDNA BR Assay kit (Invitrogen, USA) using Qubit fluorometer. The DNA quality and integrity were visualized under pulsed-field gel electrophoresis at 80 volts for 9 h in 0.5x KBB buffer (51 mM Tris, 28 mM TASP, 0.08 mM EDTA, pH 8.7) (Sage Science, USA) containing SYBR Safe DNA gel staining (Invitrogen, USA).

### PacBio library preparation and sequencing

Whole genome sequencing was performed using long read PacBio RS II and SEQUEL (Pacific Biosciences, Menlo Park, USA, outsourced to NovogenAIT, Singapore). The15-kb and 20-kb SMRTbell libraries were constructed for the PacBio RSII and SEQUEL systems, respectively. For short read sequencing, the paired-end library with 150 bp was prepared and sequenced Illumina HiSeq 2000 (Illumina, San Diego, USA) that was outsourced to Novogene, USA

### Illumina library preparation and sequencing

The short paired-end library (2×150 bp) was sequenced on the Illumina instrument at Novogene (USA). These illumine reads (133X coverage) were used to correct error reads of *de novo* assembly.

### Chicago library preparation and sequencing

A Chicago library was prepared as described previously (Putnam et al., 2016). Approximately, 500ng of high molecular weight genomicDNA (mean fragment length = 60 kbp) was reconstituted into chromatin *in vitro* and fixed with formaldehyde. Fixed chromatin was digested with DpnII, the 5’ overhangs filled in with biotinylated nucleotides, and then free blunt ends were ligated. After ligation, crosslinks were reversed to remove protein from DNA. Purified DNA was treated to remove biotin that was not internal to ligated fragments. The DNA was then sheared to ~350 bp mean fragment size and sequencing libraries were generated using NEBNext Ultra enzymes and Illumina-compatible adapters. Biotin-containing fragments were isolated using streptavidin beads before PCR enrichment of each library. The libraries were sequenced on an Illumina HiSeq X to produce 444 million 2×150 bp paired end reads, which provided 51.89 x physical coverage of the genome (1-100 kb pairs).

### Dovetail Hi-C library preparation and sequencing

A Dovetail Hi-C library was prepared in a similar way as described previously (Lieberman-Aiden et al., 2009). For each library, chromatin was fixed in place with formaldehyde in the nucleus, and then extracted fixed chromatin was digested with DpnII, the 5’ overhangs filled in with biotinylated nucleotides, and then free blunt ends were ligated. After ligation, crosslinks were reversed to remove protein from DNA. The purified DNA were then processed as similar as aforementioned in Chicago library preparation. The libraries were sequenced on an Illumina HiSeq X to produce 430 million 2×150 bp paired end reads, which provided 24,424.63X physical coverage of the genome (10-10,000 kb pairs).

### Genome assembly and scaffolding

Three types of sequencing reads from Illumina, PacBio, and Dovetail (Chicago and Hi-C reads) were used to construct *P*. *monodon* genome. PacBio sequence data was used for *de novo* assembly and Illumina sequence data was subsequently used for polishing to obtain high quality contigs. Chicago and Hi-C reads were used for scaffolding on the high quality contigs to obtain high quality *P*. *monodon* draft genome. In brief, high quality Illumina reads were prepared using TrimGalore (https://github.com/FelixKrueger/TrimGalore) based on the following criteria: (i) no “N” base, (ii) trimming of adaptor sequences and low quality bases (Q<20), (iii) no trimmed reads < 100 bp. To avoid mis-assembly due to repetitive sequences (Tørresen et al., 2019), PacBio SEQUEL subreads with repetitive sequences comprised over 85% of total sequences were filtered out. The GC content criteria (<25% and >85%) was applied for filtering low complexity DNA sequences before assembly. Noted that GC content of crustacean are 35-41% (Gao et al., 2017; Yu et al., 2015; Zhao et al., 2012). Moreover, the reads matched mitochondria sequence (NC_002184.1) were excluded from nucleus sequences and processed separately (Supplemental method). The reads ≥ 5,000 bp were assembled using WTDBG2 (Hu et al., 2019). Illumina reads were then aligned to the assembled contigs by minimap2 (Li, 2018) and subjected for polishing using wtpoa-cns mode in WTDBG2 (Hu et al., 2019).

Scaffolding of the genome assemblies was performed using HiRise, a software pipeline designed specifically for using proximity ligation data to scaffold genome assemblies (Putnam et al., 2016). The input *de novo* assembly, shotgun reads, Chicago library reads, and Dovetail Hi-C library reads were used as input data for HiRise. An iterative analysis was conducted. First, shotgun and Chicago library sequences were aligned to the draft input assembly using a modified SNAP read mapper (http://snap.cs.berkeley.edu). The separations of Chicago read pairs mapped within draft scaffolds were analyzed by HiRise to produce a likelihood model for genomic distance between read pairs, and the model was used to identify and break putative misjoins, to score prospective joins, and make joins above a threshold. After aligning and scaffolding Chicago data, Dovetail Hi-C library sequences were aligned and scaffolded following the same method. After scaffolding, shotgun sequences were used to close gaps between contigs.

The *P. monodon* genome sequence was aligned to the Pacific white shrimp *L. vannamei* (Zhang et al., 2019) using Mugsy v1.2.3 (Angiuoli & Salzberg, 2011). Alignments with a length <◻1 kb were filtered out. The output alignments between genomes were visualized using Circos v0.69◻9 (Krzywinski et al., 2009).

### Repetitive element analysis

Species-specific repeat library was generated using RepeatModeler2 (Flynn et al., 2019) prior masking with RepeatMasker (Smit, Hubley, & Green, 2013-2015). Annotation of repeats was aligned to Repbase using RMBlast. All processes were performed with default parameters.

### RNA isolation, PacBio Iso-Seq library preparation sequencing and analysis

Total RNA was extracted from gill, heart, hemocyte, hepatopancreas, intestine, ovary, testis, pleopods, stomach and thoracic ganglia of 4-month-old shrimps using TRI REAGENT according to the manufacturer’s instruction (Molecular Research Center, USA). Contaminated genomic DNA was removed by treatment with DNase I at 0.5 U/μg total RNA at 37 μC for 30 min. The DNA-free RNA was subjected for sequencing analysis using PacBio Iso-Seq SEQUEL platform (Pacific Biosciences, Menlo Park, USA, outsourced to NovogenAIT, Singapore) and ONT platform. Sequences obtained from Iso-Seq were prepared as described in Pootakham study (Pootakham et al., 2020).

### Gene prediction and annotation

Gene prediction and protein-coding sequence identification were performed using a combination of transcriptome-based prediction, homology-based prediction and *ab initio* prediction methods using EvidenceModeler (Haas et al., 2008) to generate consensus gene prediction for training species-specific parameter in AUGUSTUS (Stanke, Diekhans, Baertsch, & Haussler, 2008). To locate intron and exon regions, transcriptome-based prediction methods combined information from PacBio Iso-seq and other available *P. monodon* transcriptome databases (PRJNA4214000, SRR1648423, SRR1648424, SRR2191764, SRR2643301, SRR2643302, SRR2643304, SRR2643305) (Supporting Information Table S1) to align against the genome sequence. For short-read transcripts, STAR (Dobin et al., 2013) was employed to align against the genome before spliced read information was generated according to a previously published protocol (Hoff & Stanke, 2019) using bam2wig script in AUGUSTUS. Iso-Seq raw reads containing both 5′ and 3′ adapters (derived from full-length transcripts) were identified, and the adapters and poly(A) sequences were trimmed. Cleaned consensus reads were then mapped using Genomic Mapping and Alignment Program (GMAP) (Wu & Watanabe, 2005). Expressed sequence tags (Anchalee Tassanakajon et al., 2006) were aligned to the genome using BLAT (Kent, 2002) and converted to potential gene structures using blat2hints script of AUGUSTUS.

Protein sequences of *P. monodon* were mapped against proteins from closely related organisms including *H. azteca*, and *L. vannamei*, using Exonerate version 2.2.0 (Slater & Birney, 2005). All gene models derived from these three methods were integrated by EvidenceModeler into a high confident nonredundant gene set, which was used to set species-specific parameters. Finally, AUGUSTUS was used to predict genes in the genome based on the extrinsic evidence. Functional annotations of the obtained gene set were conducted using Semantic Annotation Platform (SAPP) using the InterProScan module (P. Jones et al., 2014; Koehorst et al., 2018) and Blast2GO (Götz et al., 2008).

### Phylogenetic analysis

Mitochondria protein-coding genes were used to construct a molecular phylogenetic analysis using MEGA7 software (Kumar, Stecher, & Tamura, 2016). The amino acid sequences of 22 species, which were *Anopheles quadrimaculatus*, *Anopheles gambiae*, *Apis mellifera,, Artemia franciscana, Ceratitis capitata, Daphnia pulex*, *Drosophila yakuba*, *Drosophila melanogaster*, *Euthynnus* affinis, *Halocaridina rubra*, *Hyalella azteca, Ixodes hexagonus*,, *L. vannamei*, *Ligia oceanica*, *Locusta migratoria*, *Macrobrachium rosenbergii*, *Panulirus japonicus*, *Parhyale hawaiensis, P. monodon*, *Portunus trituberculatus*, *Rhipicephalus sanguineus*, and *Tigriopus japonicus* were used for the analysis. The evolutionary history was inferred by using the maximum likelihood method based on the JTT matrix-based model (D. T. Jones, Taylor, & Thornton, 1992). The tree with the highest log likelihood (−84614.94) is shown. An initial tree for heuristic search was obtained automatically by applying Neighbor-Join and BioNJ algorithms to a matrix of pairwise distances estimated using a JTT model, and then selecting the topology with superior log likelihood value.

Pan-core protein families were constructed for *P. monodon* with the available crustacean genome *E. affanis*, *D. pulex*, *P. hawaiensis*, *L. vannamei* and *H. azteca.* The orthologous genes protein family were identified by clustering together using MMseq2 (Steinegger & Söding, 2017) with standard parameters of 0.5 identity and 0.5 coverage. The pan-core protein family matrix was used to construct a pan-core protein family tree follow the method presented by Snipen et al (Snipen & Ussery, 2010). The comparison of common and specific protein family among the crustacean genomes was present in the upset plot (Lex, Gehlenborg, Strobelt, Vuillemot, & Pfister, 2014). The pan-core protein family tree was used to construct an ultrametric tree using ape v5.3 library in R v3.6.1 (Paradis & Schliep, 2018; R Core Team, 2019). The ultrametric tree and matrix of protein families were subjected to CAFE v4.1 (Bie, Cristianini, Demuth, & Hahn, 2006) to study the evolution of gene families in *P. monodon*. The resulting families were further filtered to exclude families with large ranges in size (> 25) across the tree, leaving 102619 families for expansion and contraction analysis. CAFE was used to obtain a maximum likelihood estimate of a global birth and death rate parameter λ (lambda; the rate of gain/loss per gene per million years) across the species tree (λ =0.27). The size of families was estimated at each ancestral node and then used to obtain a family-wise p-value to indicate non-random expansion or contraction for each family and across all branches of the tree.

### Transcriptome sequencing and analyses of the fast-growing and slow-growing groups

The feeding trial was carried out at SGIC. Black tiger shrimps were cultured in a pond size 800 m^2^ (25 shrimps/m^2^). Five-month-old female shrimps (*N*=140) were randomly selected for body weight measurement (Supporting Information Figure S1). Two experimental groups were separated base on the lowest and highest shrimp weight, called the “slow-growing” group and the “fast-growing” group, respectively. Average body weight of 15 samples from the lowest body weight of the slow-growing group was 13.46±0.52 g, while the average body weight of 15 samples from the highest of the fast-growing group was 36.27±1.96 g (Supporting Information Table S2). All hepatopancreas samples were immediately frozen in liquid nitrogen and stored at −80°C until use. To extract total RNA, individual frozen hepatopancreas tissues were pulverized in liquid nitrogen and subjected to TRI-REAGENT extraction method and DNase treatment as previously described.

To obtain short-read RNA sequences, 30 libraries (n=15 for each group) were constructed using the HiSeq Library Preparation kit (Illumina, San Diego, USA) and sequenced using Illumina NovaSeq™. Illumina sequencing using Novaseq 6000 instrument with 150 pair-end reads was performed at Omics Drive, Singapore. The raw reads were quality-filtered (Q<20, >50bp) and adapter-trimmed using TrimGalore (https://github.com/FelixKrueger/TrimGalore). The reads were mapped to the genome using STAR (Dobin et al., 2013). The HTSeq-count(Anders, Pyl, & Huber, 2015) and DESeq2 (Love, Huber, & Anders, 2014) provide a test for differential expression using negative binomial generalized linear models, will operate to identify significant differently expressed genes (DEGs) between fast- and slow-growing shrimp. DEGs were identified when their expression level differences > 2.0 change with Bonferroni adjusted p-value of < 0.05. The DEGs which have read count per million < 1 were discarded. Functional pathway analysis was carried using EggNOG (Huerta-Cepas et al., 2019) and KEGG mapper (Kanehisa, Sato, Kawashima, Furumichi, & Tanabe, 2016). To determine whether the differentially expressed genes may be newly identified genes in *P. monodon*, the sequences were aligned on available *P. monodon* protein and mRNA sequences in NCBI database using BLAST (E-value ≤ 10^−3^). The sequences that had no match to any publicly available *P. monodon* sequences but could be annotated were considered newly identified genes.

To evaluate differentially expressed genes, 15 genes (10 DEGs that had higher levels of expression in the fast-growing shrimp and 5 DEGs that had higher levels of expression in the slow-growing shrimp) were selected for real-time PCR analysis. Eight hepatopancreas per group (fast-growing and slow-glowing) were used for validation. Total RNA (1.5 μg) was reverse transcribed into cDNA using an ImPromII™ Reverse Transcriptase System kit (Promega, USA.) according to the manufacturer’s recommendation. Each 20 μL qPCR reaction included 200 ng cDNA, 0.2 μM of each primer, and SsoAdvanced™ Universal SYBR^®^ Green supermix (BioRad, USA.) according to the company’s instruction. The thermal cycling parameters were 95°C for 2 min 30 s, followed by 40 cycles of 95°C for 15 s, 58°C for 20 s and 72°C for 30 s. The melting curve analysis was performed from 65°C to 95°C with a continuous fluorescent reading with a 0.5°C increment. The threshold cycle (Ct) was analyzed using BioRad CFX Manager 2.1 software (BioRad, USA).

## Results

### Genome sequencing, assembly and annotation

A whole-genome shotgun strategy was used to sequence and assemble a black tiger shrimp genome from PacBio long-read data. A total of 13,157,113 raw reads (178.94 Gb) representing 69.08X coverage based on the estimated genome size of 2.59 Gb obtained from a previous report using the flow cytometry method (Swathi, Shekhar, Katneni, & Vijayan, 2018). *De novo* assembly of PacBio sequences yielded a preliminary assembly of 2.39 Gb (70,380 contigs) with a contig N50 of 79 kb and a L50 of 6,786 contigs (Table 1). The draft genome was further assembled using the long-range Chicago (*in vitro* proximity ligation; 444 million read pairs) and Hi-C (*in vivo* fixation of chromosomes; 430 million read pairs) library data scaffolded with the HiRise software (Dovetail Genomics, Santa Cruz, USA). The final assembly contained 44 pseudomolecules greater than 5 Mb in length (hereafter referred to as chromosomes, numbered according to size; Figure 1A), corresponding to the haploid chromosome number in *P. monodon* (1*n* = 44, 2*n* = 88). The 44 chromosomes covered 1,986,035,066 bases or 82.9% of the 2.39-Gb assembly.

**Table 1.**
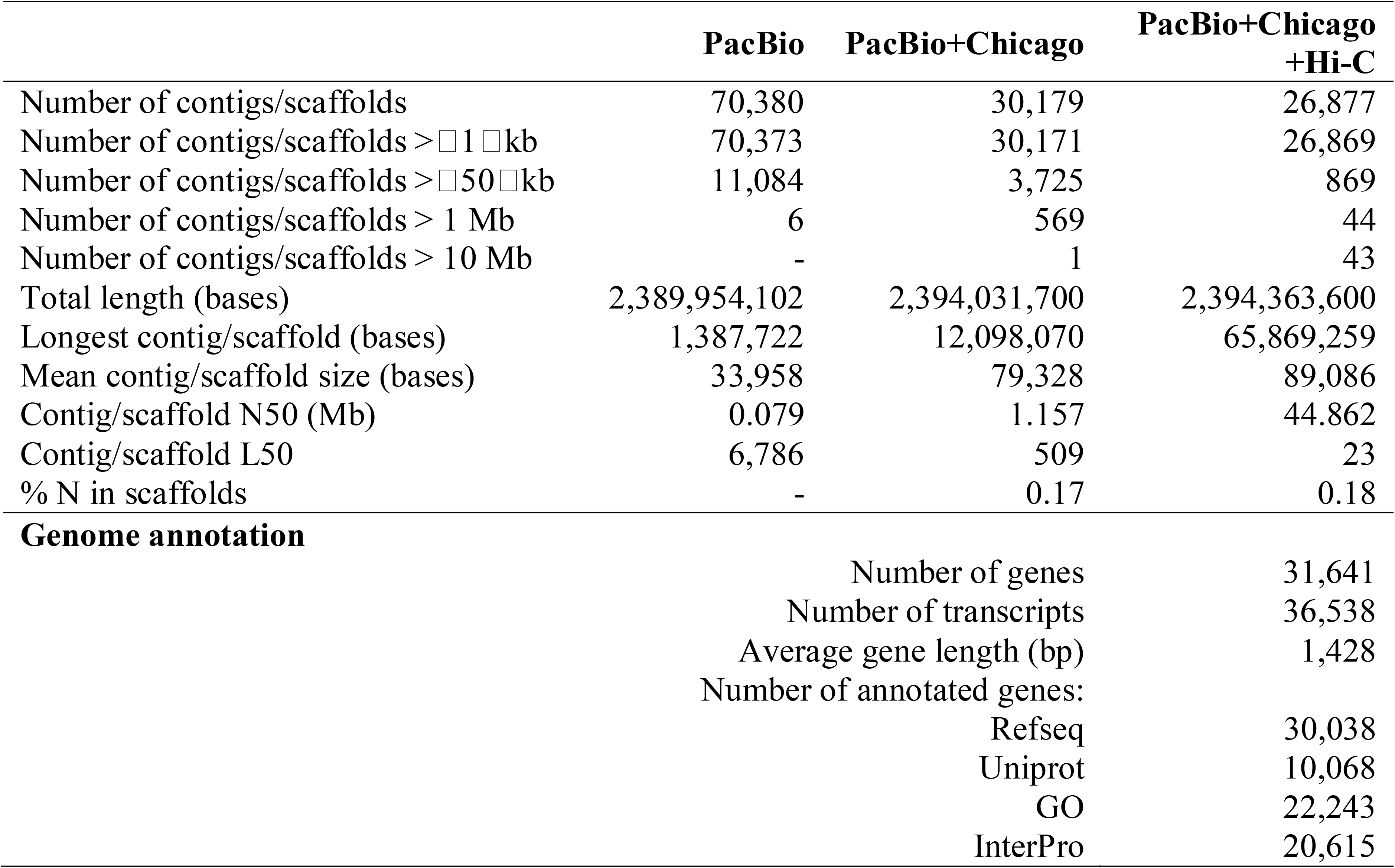
Assembly statistics of the *P. monodon* genome.

|  | PacBio | PacBio+Chicago | PacBio+Chicago<br>+Hi-C |
| --- | --- | --- | --- |
| Number of contigs/scaffolds | 70,380 | 30,179 | 26,877 |
| Number of contigs/scaffolds > 1 kb | 70,373 | 30,171 | 26,869 |
| Number of contigs/scaffolds > 50 kb | 11,084 | 3,725 | 869 |
| Number of contigs/scaffolds > 1 Mb | 6 | 569 | 44 |
| Number of contigs/scaffolds > 10 Mb | - | 1 | 43 |
| Total length (bases) | 2,389,954,102 | 2,394,031,700 | 2,394,363,600 |
| Longest contig/scaffold (bases) | 1,387,722 | 12,098,070 | 65,869,259 |
| Mean contig/scaffold size (bases) | 33,958 | 79,328 | 89,086 |
| Contig/scaffold N50 (Mb) | 0.079 | 1.157 | 44.862 |
| Contig/scaffold L50 | 6,786 | 509 | 23 |
| % N in scaffolds | - | 0.17 | 0.18 |
| **Genome annotation** |  |  |  |
|  | Number of genes |  | 31,641 |
|  | Number of transcripts |  | 36,538 |
|  | Average gene length (bp) |  | 1,428 |
|  | Number of annotated genes: |  |  |
|  | Refseq |  | 30,038 |
|  | Uniprot |  | 10,068 |
|  | GO |  | 22,243 |
|  | InterPro |  | 20,615 |

**Figure 1.**
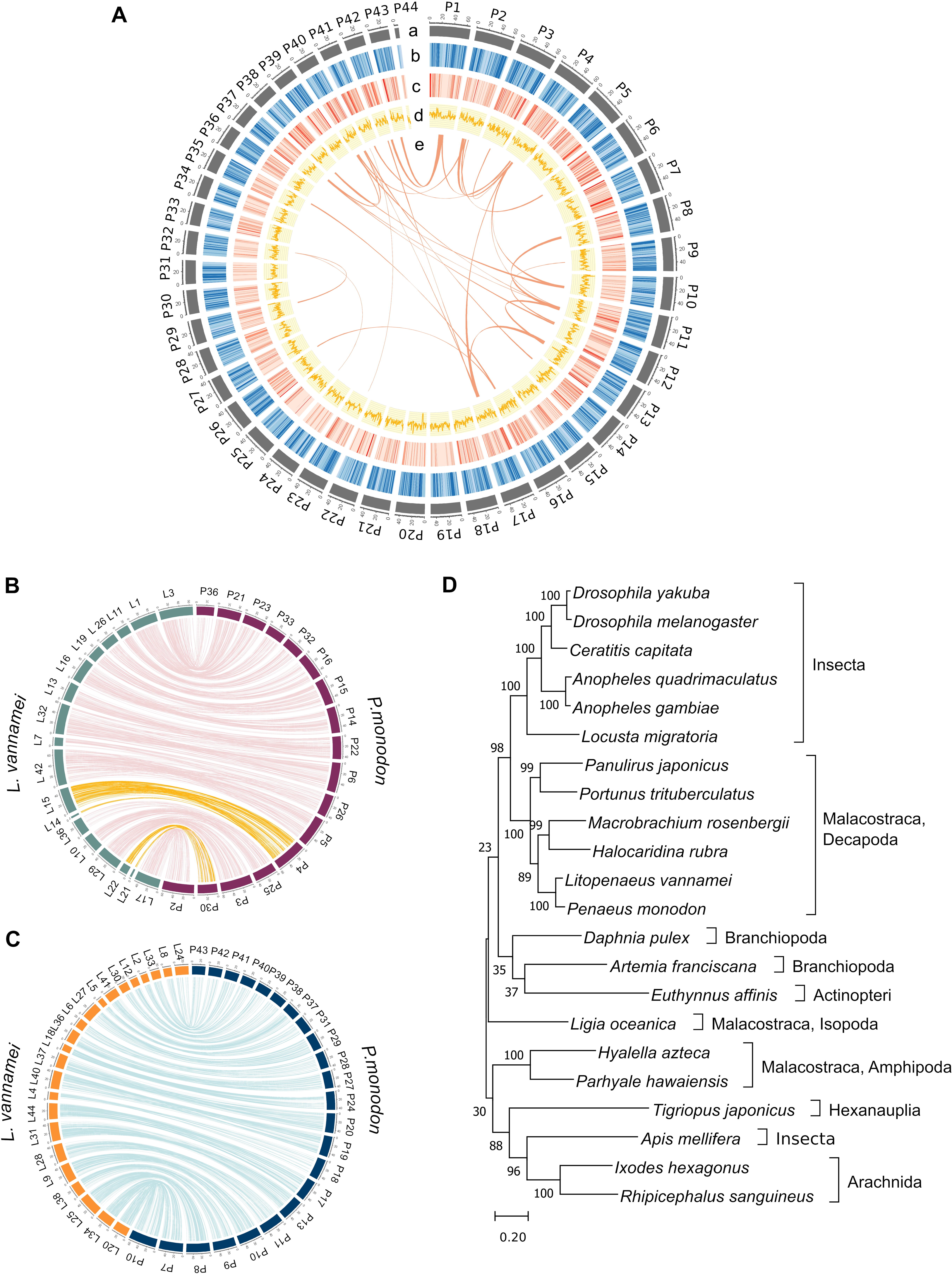
*P. monodon* genome assembly and phylogenetic analysis. (**A**) Genomic landscape of 44 assembled *P. monodon* chromosomes. (a) Physical map of *P. monodon* chromosomes (Mb scale). (b) Density of repetitive sequences represented by percentage of genomic regions covered by simple repeat sequences in 500-kb window. (c) Gene density represented by number of genes in 500-kb window. (d) GC content represented by percentage of G/C bases in 500-kb window. Syntenic blocks are depicted by connected lines. (e) Syntenic relationship of gene blocks among *P. monodon* chromosomes. Syntenic blocks were identified by MCScanX with criteria at least ten syntenic genes and a maximum of six intervening genes allowed. (**B**) Diagrams showing colinearity between *P. monodon* and *L. vannamei* chromosomes. Lines link the position of orthologous gene sets. Most regions exhibit one-to-one relationship between *P. monodon* and *L. vannamei.* The yellow line represents the region on a single *P. monodon* chromosome that exhibits synteny to regions on two *L. vannamei* chromosomes. Black tiger shrimp chromosomes are designated with “P” followed by chromosome numbers, and Pacific white shrimp chromosomes are designated with “L” followed by the chromosome numbers. (**C**) Diagrams showing colinearity between *P. monodon* and *L. vannamei* where the syntenic relationship between chromosomes is not one-to-one. (**D**) Phylogenetic tree of concatenated mitochondria protein-coding genes using maximum likelihood general reversible mitochondrial model. The percentage of trees in which the associated taxa clustered together is shown next to the branches. The tree is drawn to scale, with branch lengths measured in the number of substitutions per site.

To evaluate the quality of our *de novo* assembly, we aligned DNA short reads obtained from Illumina sequencing from this work and the previous report (Yuan et al., 2018) to the assembled genome and found that approximately 93% of the DNA short reads could be mapped on the here reported *de novo* assembly. We also aligned the publicly available RNA-seq reads (Huerlimann et al., 2018) and Iso-seq reads (Pootakham et al., 2020) to the assembly, and 90.22% and 98.77% of the RNA-seq and Iso-seq reads were mapped to the assembly, respectively. To further assess the completeness of the genome assembly, we checked the gene content with the BUSCO software using the Eukaryota (odb9) database (Simão, Waterhouse, Ioannidis, Kriventseva, & Zdobnov, 2015). Our gene prediction contained 94.72% of the highly conserved orthologs (85.15% complete, 9.57% partial, 5.28% missing) in the eukaryotic lineage. High mapped rates, comparable to the numbers observed in *L. vannamei*, and a high percentage of identified highly conserved orthologs provided the evidence for a high-quality assembly obtained for the black tiger shrimp genome.

The chromosome-level contiguity achieved in our assembly enabled a syntenic analysis between *P. monodon* and *L. vannamei*. Portions of conserved syntenic genes between the black tiger shrimp and Pacific white shrimp are illustrated in Figure 1B and 1C. The degree of macrosynteny observed between these two species was consistent with the result from phylogenetic analysis showing that they are very closely related and appear to have evolved from a common ancestor (Figure 1D). The distribution of paralogous gene pairs revealed one-to-one synteny across 15 chromosomes/linkage groups and one-to-two synteny between *P. monodon* chromosomes 4 and 30 and *L. vannamei* chromosomes 14, 15, 21 and 22 (Figure 1B).

A combination of *ab initio* prediction, homology-based search and transcript evidence obtained from both Iso-seq and RNA-seq data was used for gene prediction. The genome annotations of *P. monodon* contained 31,640 predicted gene models, of which 30,038 were protein-coding genes (Table 1, Supporting Information Table S3). The number of protein-coding genes in *P. monodon* was slightly higher than those reported in *L. vannamei* (25,596) (Zhang et al., 2019) and in a freshwater epibenthic amphipod *Hyalella azteca* (19,936) (Poynton et al. 2018). Of 30,038 protein-coding genes, 25,569 (85.02%) had evidence support from our RNA-seq and Iso-seq data.

In addition to the nuclear genome, a complete mitochondrial genome assembly was obtained, enabling phylogenic analysis to reveal relationship with other arthropods. The assembled mitochondrial genome had a size of 15,974 bp and 29.09% GC content (Supporting Information Figure S2). The phylogenetic analysis of 20 arthropods based on 13 concatenated mitochondrial protein-coding genes showed that *P. monodon* and other Decapoda were more closely related to the insects than other crustaceans in the same class (Malocostraca) such as *H. azteca* and *Parhyale hawaiensis* (Amphipoda), consistent with a previous study (Wilson, Cahill, Ballment, & Benzie, 2000) (Figure 1D). The phylogenic tree also showed that decapods were distantly related to crustaceans in Branchiopoda class such as *Artemia franciscana* and *Daphnia pulex.*

### Comparative analyses between *P. monodon* and other crustacean genomes

Comparative analyses of *P. monodon* genome with publicly available crustacean genomes (Calanoida (*Eurytemora affinis*) (Eyun et al., 2017), Cladocera (*D. pulex*) (Colbourne et al., 2011) and Amphipoda (*P. hawaiensis* (Kao et al., 2016)*, L. vannamei* (Zhang et al., 2019) and *H. azteca* (Poynton et al., 2018)) were performed to investigate genome evolution within the crustacean group. Based on orthologous clustering by MMseq2 software (Steinegger & Söding, 2017), we identified 102,632 pan protein families of the crustacean with the core protein family of 583 (Figure 2A). The dendrogram analysis of protein families showed a distinct group of the *Amphipoda*. As expected of the closely related species, *P. monodon* and *L. vannamei* shared 5,487 common protein families that were higher than those between *P. monodon* and other species. Nevertheless, both species also had around 11,000 species-specific protein families (Figure 2A). The numbers of expanded and contracted gene numbers at each divergence event were estimated using CAFÉ software (Bie et al., 2006). Considerable gene family expansions occurred on the *P. monodon* branch that might have experienced additional expansions after its divergence from *H. azteca.*

**Figure 2.**
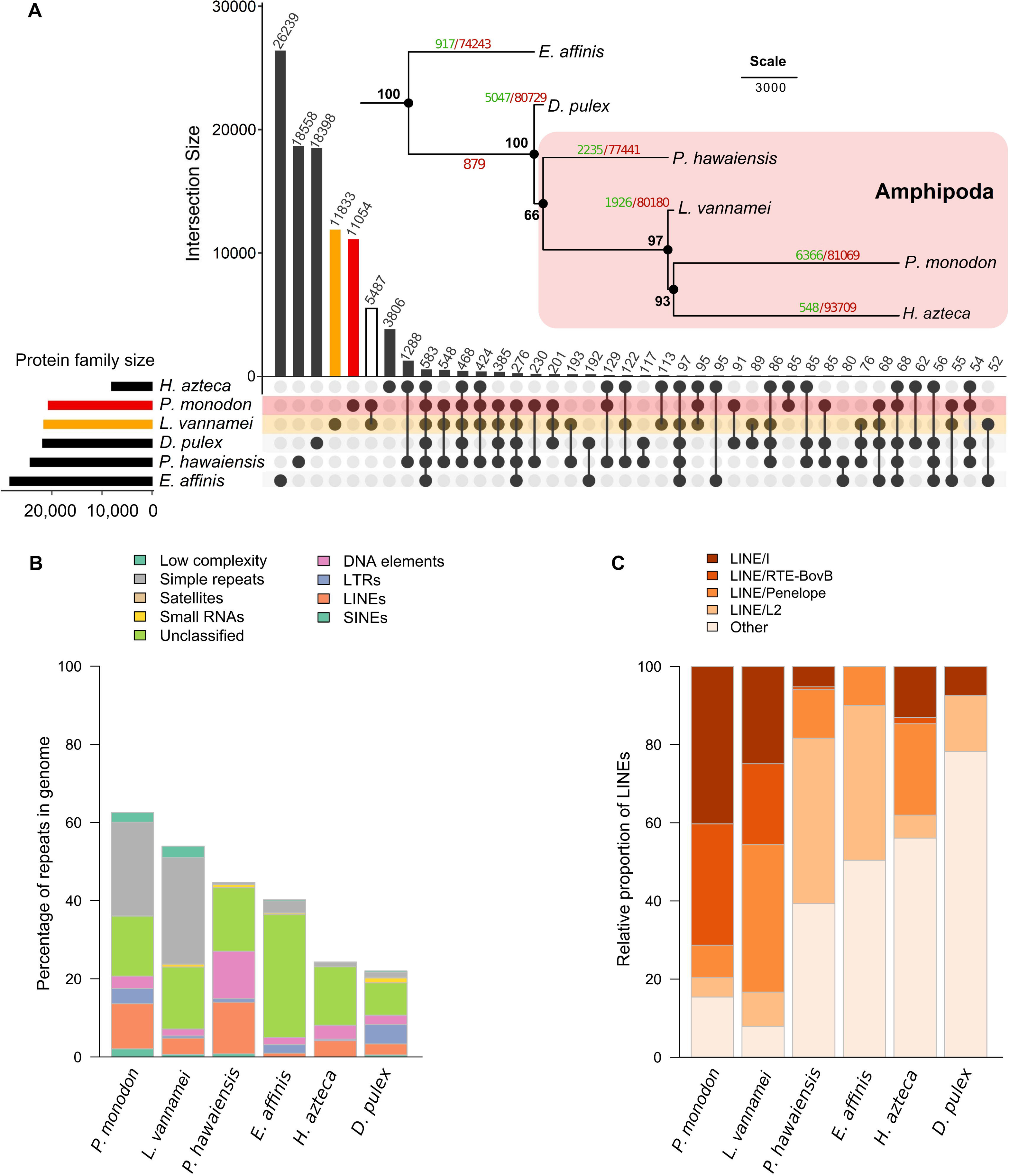
Comparative genomics and repeat elements analysis. (**A**) Upset plot represents pan-core protein family of crustacean genomes (*P. monodon, L. vannamei, P. hawaiensis, E. affinis, H. azteca,* and *D. pulex*) with the pan protein family tree with bootstrap value in black numbers. The numbers on branches indicate the number of protein family gains (green) or losses (red). Relative proportion or repeat elements in the crustacean genomes is presented as (**B**) bar plots of percentage of repeat elements and (**C**) relative abundance of each LINE type.

### Repetitive elements in *P. monodon* genome

The total repeat content in the black tiger shrimp genome assembly was 62.5% (Figure 2B). We identified repeats comprising 572.87 Mb simple repeats (23.93%), 276.64 Mb long interspersed nuclear elements (LINEs, 11.55%), 93.21 Mb long terminal repeats (LTRs, 3.89%), 75.96 Mb DNA elements (3.17%), 59.09 Mb low complexity repeats (2.47%), 49.49 Mb short interspersed nuclear elements (SINEs, 2.07%), 2.18 Mb small RNA (0.09%), and 368.13 Mb of unclassified repeat elements (15.37%).

Simple repeats and LINEs are the two most abundant repeat categories in *P. monodon,* together accounting for 35.48% of the genome assembly. Moreover, LINE/I (4.65%), RTE-BovB (3.59%), and Penelope (0.96%) were the three major components of LINEs (Figure 2C). The major LINEs components found here were in accord with the previously reported *P. monodon* genome, but with different proportions (2.03%, 4.96% and 0.82% for LINE-I, RTE-BovB, and Penelope, respectively) (Yuan et al., 2018).

### Identification of novel growth-associated genes

Transcriptome analysis is a useful approach to identify genes that are differentially expressed among individuals with various traits of interest. Without a high-quality reference genome, gene expression studies have to rely on a *de novo* transcriptome assembly, which often contains a large number of fragmented transcript sequences with no annotation or in some cases incorrect annotations. To evaluate the utility of this high-quality genome assembly in transcriptomic analysis, transcriptomics of hepatopancreas from the fast- and slow-growing shrimps were compared with and without the genome assembly as a reference.

Comparison of the overall results from the genome assembly as a reference and from *de novo* transcriptome assembly revealed a significant improvement. With the genome assembly, a lower number of predicted genes with a higher N50 value was obtained (31,640 genes with N50 of 1,743 nt from the genome assembly vs 340,240 genes with N50 of 776 nt from the *de novo* transcriptome assembly). Moreover, a higher annotation rate (95.01%) was obtained with the genome assembly as a reference than that from the *de novo* transcriptome assembly (26.36%).

We compared the differential gene expression profile in hepatopancreas of the fast-growing shrimp and the slow-growing shrimp. This analysis identified 383 genes exhibiting higher levels of expression in the fast-growing shrimp and 95 genes exhibiting higher levels in the slow-growing shrimp (Log_2_ fold-change>1 and *p*-value <0.05) (Supporting Information Table S4). The fast-growing shrimp grew at a faster rate and had twice the weight of the slow-growing shrimp under the same rearing condition at 5 months old. To further access gene interaction networks, KEGG pathway mapping was employed on the DEGs and revealed 159 pathways with an average of two genes associated in each pathway (Supporting Information Table S5). The functions of DEGs were classified by Clusters of Orthologous Groups (COGs) annotation into 23 categories (Supporting Information Figure S3). The top five highly enriched metabolic processes were carbohydrate/lipid/amino acid transport and metabolism, secondary metabolites biosynthesis, transport and catabolism, and inorganic ion transport and metabolism (Figure 3A). For instance, DEGs involved in carbohydrate metabolism were *amylase* (*amy*), *fructose-bisphosphate aldolase* (*aldo*), *glyceraldehyde 3-phosphate dehydrogenase* (*gapdh*), and *insulin-like growth factor-1 receptor* (*insr*). Genes involved in lipid metabolism were *nose resistant to fluoxetine* (*nrf*), *lipase* (*lip*), g*lycerol*-*3*-*phosphate acyltransferase* (*gpat3*)*, acyl-CoA delta desaturase* (*scd*)*, long-chain-fatty-acid--CoA ligase* (*acsl1*) and *elongation of very-long-chain fatty acids protein 4-like* (*elovl4*). Of all DEGs, 67 annotated genes could not be matched with any *P. monodon* sequences in the publicly available genomic/transcriptomic databases (Supporting Information Table S4). These newly identified genes involved in nutrient metabolic processes (Figure 3A shown in blue). We further investigated pathways that have not been reported to be involved in shrimp growth and found that PI3K-Akt signaling pathway has the highest number of novel genes (Figure 3B). Four DEGs in PI3K-Akt signaling pathway were *integrin beta isoform 1* (*itgb1*), *integrin alpha-4 like* (*itga4*), *serine/threonine-protein kinase N isoform 1* (*pkn1*) and *insr.* Of these, the last three were newly identified genes.

**Figure 3.**
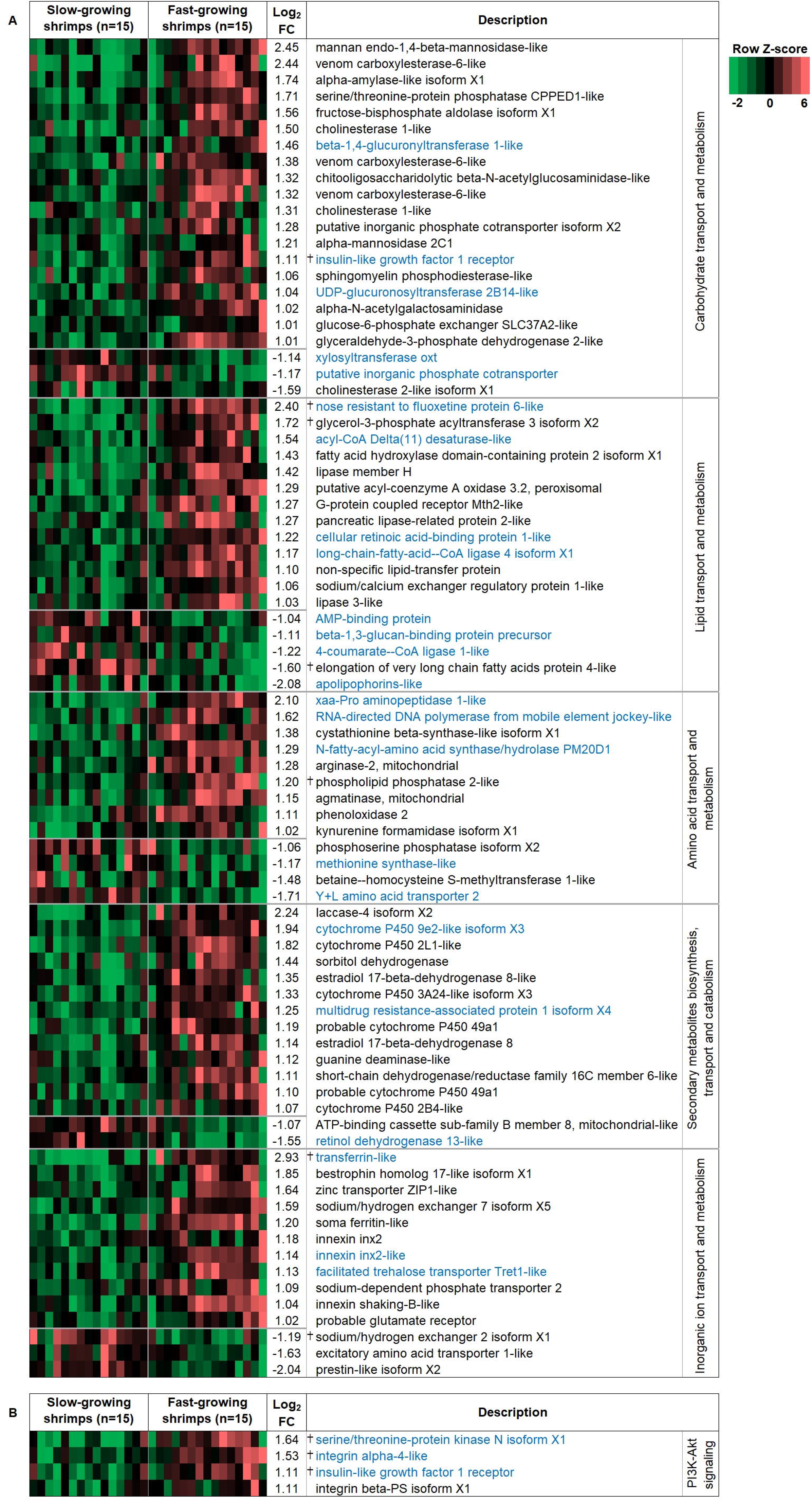
Expression of differentially expressed genes of slow-growing shrimp (n=15) and fast-growing shrimp (n=15). (**A**) The top five enriched COGs metabolism groups were carbohydrate transport and metabolism, secondary metabolites biosynthesis, transport and catabolism, inorganic ion transport and metabolism, lipid transport and metabolism and amino acid transport and metabolism. (**B**) Differentially expressed genes in PI3K-Akt pathway. Blue letters indicate newly identified genes in *P. monodon*, and the dagger (†) indicates genes that were further validated by quantitative real-time PCR.

To verify the transcriptome analysis, 15 DEGs with functions related to nutrient metabolism or immune system were selected for quantitative real-time PCR (qPCR). The qPCR results agreed with those obtained from RNA-seq with a correlation coefficient of 0.96 (Supporting Information Figure S4).

## Discussion

In this study, we present a whole genome sequence of *P. monodon* assembled from PacBio long-read data and scaffolded using long-range Chicago and Hi-C techniques. Our chromosome-scale assembly has shown tremendous improvement in contiguity and completeness compared to the previously reported *P. monodon* genomes (Quyen et al., 2020; Yuan et al., 2018). Based on a benchmark (10Mb) (Reference standard for genome biology, 2018), the assembly presented here is considered a high-quality reference genome as it has the N50 scaffold length of 44.86 Mb. It is also one of the highest quality crustacean genomes currently available. Of the 45 crustacean genome sequences listed in the NCBI genome database, only three species (*Tigriopus japonicus*, *T. californicus* and *Daphnia magna*) have their genomes assembled at a chromosome level.

The present genome provides an invaluable resource for shrimp research. The availability of the chromosome-scale assembly allowed us to examine the syntenic relationship between the two economically important penaeid shrimp species: the black tiger shrimp and the Pacific white shrimp genomes. The synteny analysis revealed a one-to-one relationship between *P. monodon* and *L. vannamei* chromosomes, suggesting that certain chromosomes derived from the common ancestor of *P. monodon* and *L. vannamei* were fragmented into two smaller chromosomes in *L. vannamei* (for example, *P. monodon* chromosome 4 displayed synteny to *L. vannamei* pseudochromosomes 14 and 15; Figure 1B) but remained as single chromosomes in *P. monodon*.

Obtaining contiguous long-read-based genome of shrimp has been hampered by limitation related to short-read sequencing technology. PacBio sequencing reads allow the assembler to obtain a contiguous assembly that spans repeat regions containing transposable elements and tandem repeats. We found that *P. monodon* has the highest repeat abundance (62.5%) when compared to five available genome sequences of crustacean species: *L. vannamei* (53.9%), *P. hawaiensis* (44.7%), *E. affinis* (40.2%), *H. azteca* (24.3%) and *D. pulex* (22.0%). Moreover, the percentage of repeat elements observed in this assembly was substantially higher than reported in previous studies on *P. monodon* fosmid (51.8%) (Huang et al., 2011) and genome (46.8%) (Yuan et al., 2018). High repeat contents and long repetitive sequences hindergenome assembly. Ambiguous regions containing mostly repetitive sequences might be missed or caused errors in assembly using short sequencing reads employed in the previous assembly of the *P. monodon* genomes (Yuan et al., 2018) and *L. vannamei* genome (Zhang et al., 2019). Our assembly, on the other hand, utilized the PacBio sequencing platform, which yielded kilobase-sized reads that could be assembled into contigs and scaffolds large enough to span repetitive regions, alleviating the problems often encountered by the use of short-read technologies.

The chromosome-scale assembly allows for an in-depth investigation of repeat elements. Even though the biological function of repeats has not been well studied in the shrimp, they might be associated with important functions as reported in other organisms. For example, LINE/I plays a role on transcription in human by co-mobilizing DNA to new locations (Pickeral, Makałowski, Boguski, & Boeke, 2000). Among diverse repetitive elements, LINE was the major element in both *P. monodon* and *L. vannamei*. The most abundant element in *P. monodon* and *L. vannamei* were LINE/I and LINE/L2, respectively. Given that the diversity of repeats was deemed to play a role in environmental adaptation of animals (Schrader & Schmitz, 2019), roles of these repetitive elements in the black tiger shrimp could be further explored to gain a better understanding of shrimp biology.

Another advantage of the high-quality reference genome is the reduction in erroneous and fragmented assembled contigs using *de novo* transcripts. With this high-quality reference genome assembly, we were able to obtain an improved gene set with better contiguity and annotation rate. Of the predicted genes in the genome, 95.01% of them could be functionally annotated. The results suggested that our genome assembly could serve as a high-quality reference for facilitating functional genomic study in the black tiger shrimp.

The chromosome-scale assembly facilitates downstream applications for molecular breeding and gene expression studies in shrimp. Growth is undoubtedly an important factor for profitable shrimp production. However, a daunting challenge in black tiger shrimp farming is its slower growth rate in captivity (Benzie, Kenway, & Ballment, 2001; Cheng & Chen, 1990). Domesticated shrimps could not mature well with declining growth rates over generations (Jackson & Wang, 1998). Albeit its importance, the knowledge on genes controlling shrimp’s growth remained limited partly due to the lack of the genome, thus up-to-now most growth-related genes identified in *P. monodon* were from *L. vannamei* (Gao et al., 2017; Gao et al., 2015; Santos et al., 2018).

The comparison of hepatopancreas transcriptomes of slow-growing shrimp and fast-growing shrimp revealed that DEGs were mainly involved in nutrient metabolism, which was in concordance with the hepatopancreas functions in feed utilization and energy storage. Here, we presented genes related to carbohydrate and lipid metabolisms as they are main nutrients that have been investigated to enhance growth in shrimp (Coelho, Yasumaru, Passos, Gomes, & Lemos, 2019; González-Félix, Gatlin, Lawrence, & Perez-Velazquez, 2002; Hu et al., 2019; Olmos, Ochoa, Paniagua-Michel, & Contreras, 2011). Genes with higher expression levels in the fast-growing group were found to be involved in nutrient metabolism and secondary metabolite biosynthesis. Genes exhibiting higher levels of expression in the fast-growing shrimp were digestive enzymes such as amylase and lipase, in agreement with the prior finding that enhancement of digestive enzyme activities improves growth performance of shrimp activities (Anand et al., 2013; Gómez & Shen, 2008). As dietary carbohydrates can enhance growth (Sagar et al., 2019), it is not surprising to find carbohydrate metabolism genes such as *aldo*, *gapdh* and *insr* expressed at a higher level in the fast-growing shrimp

Similarly, DEGs in lipid metabolism agree with the established knowledge on the importance of fatty acids and lipids to the shrimp growth and immunity (Chen et al., 2015; Duan et al., 2019; Toledo, Silva, Vieira, Mouriño, & Seiffert, 2016). Many fatty acids such as highly unsaturated fatty acids (HUFA) are indeed essential to marine animals since they are the major component of cell membrane (An et al., 2020). Thus, regulation of the synthesis, digestion and absorption of these fatty acids is a key to shrimp growth. The higher expression of genes involved in unsaturated fatty acid (UFA) biosynthesis such as *scd* and *acsl1* in the fast-growing group suggested that these fatty acids might positively associate with growth. The results agree with the previous studies reported that additional of high HUFA such as linoleic (LOA, 18:2n–6), linolenic (LNA, 18:3n–3), eicosapentaenoic acid (EPA, 20:5n–3) or docosahexaenoic acid (DHA, 22:6n–3) in diet could promote the growth (Glencross, Smith, Thomas, & Williams, 2002a, 2002b). Interestingly, *elovl4* that synthesizes very long-chain (>C24) saturated and polyunsaturated fatty acids (Oboh, Navarro, Tocher, & Monroig, 2017) was expressed at a higher level in the slow-growing shrimp than in the fast-growing one. In gilthead sea bream *Sparus aurata*, high amount of LNA and long-chain fatty acid adversely affected growth (Turkmen et al., 2019). Considering the above data, it could be implied that balance lipid composition is an important factor controlling growth performance; thus, feed formulation to promote growth should be optimized by considering lipid ratios.

Among the DEGs, three new candidate marker genes may potentially be useful for shrimp breeding programs, namely *tranferin* (*trf*), *nrf* and *pkn1*. Trf is an insulin-like growth factor (IGF) that stimulates both proliferation and differentiation in a cell line (Kiepe, Ciarmatori, Hoeflich, Wolf, & ToLnshoff, 2005), making it a promising candidate as a growth marker. Nrf, a lipid-carrier protein, has been reported to be essential for embryonic development in *Caenorhabditis elegans* (Watts & Browse, 2006). Given the aforementioned importance of lipid for shrimp growth, Nrf might regulate the intake and storage of lipid for the growth. Besides nutrient metabolic pathways, PI3K-Akt pathway presents an interesting pathway for further investigation for its involvement in growth regulation as we found three novel genes (*pkn1*, *itga4*, and *insr*) in this pathway. PI3K-Akt is a regulatory pathway controlling glucose balance by cross-talking with insulin signaling pathway (Shi & He, 2016). Although an association between PI3K-Akt and shrimp growth performance has not yet been investigated in *P. monodon*, there has been a report that PI3K-Akt is linked to growth factors and cellular survival (Choi et al., 2019; Dai, Li, Fu, Qiu, & Chen, 2020; Fuentes, Valdés, Molina, & Björnsson, 2013). Here, novel genes associated with PI3K-Akt pathway showed higher expression in the fast-growing shrimp; therefore, it suggested that these genes might have potential roles in promoting growth. Particularly, the distinct expression values of *pkn1* between fast- and slow-growing shrimps make it interesting for further study (Supporting Information Figure S4). Pkn1 is found in various tissues with different functions in many animals (Mukai, 2003). For instance, Pkn1 in testis regulates male fertility of the Pacific abalone *Haliotis discus hannai*, (Kim, Kim, Park, & Nam, 2019), whereas it has been linked to the regulation of actin and cytoskeletal network in human (Dong et al., 2000). These newly identified candidate genes might provide better understanding of shrimp growth and facilitate black tiger shrimp genomic breeding programs.

In conclusion, we have successfully overcome the technical challenges in obtaining the first high-quality chromosome-scale genome assembly of the economically important *P. monodon*. The availability of this reference genome enables several downstream biological and industrial applications that would otherwise be difficult. This reference genome will benefit not only the *P. monodon* research community, but also other researchers working on related shrimp and crustacean species. Moreover, the newly identified growth-associated genes might help advance the understanding of growth in the black tiger shrimp and facilitate its genomic breeding programs.

## Acknowledgements

The authors acknowledge NSTDA Supercomputer Center (ThaiSC) for providing HPC resources that have contributed to the research results reported within this paper. We thank Ms. Somjai Wongtripop, Mr. Kaidtisak kaewlok and Shrimp Genetic Improvement Center (SGIC) members, as well as Ms. Sudtida Phuengwas (BIOTEC) for the shrimp sample collection. This project was funded by National Center for Genetic Engineering and Biotechnology (BIOTEC Platform No. P1651718), TDG Food and Feed program, National Science and Technology Development Agency (No. P1950419) and partially supported by the European Union’s Horizon 2020 research and innovation programme under the Marie Skłodowska-Curie grant agreement No 734486 (SAFE-Aqua). Visiting Professor Program 2019 was awarded to Dr. Intawat Nookaew by the National Science and Technology Development Agency, Thailand. We acknowledge Zulema Dominguez (University of Arkansas for Medical Sciences, USA) for technical assistance on comparative genome. We are grateful to Prof. Morakot Tanticharoen (NSTDA), Dr. Kanyawim Kirtikara (BIOTEC), Dr. Wonnop Visessanguan (BIOTEC) and Assoc Prof. Skorn Koonawootrittriron (Kasetsart University, Thailand) for their mentorship on the shrimp research programs.

## Data accessibility

The genomic and transcriptomic data are available in the NCBI under the following accession numbers: PRJNA611030 and PRJNA602748. The annotation file and genome sequences are available from the following website: http://www.biotec.or.th/pmonodon/index.php.

## Author contributions

PA, KS, SA, RL and JK collected samples. PA, KS, SA, TW, RL, DS extracted DNA and RNA for sequencing. TU, WP, and CS carried out genome assembly and assessment. TU, JK, PS and VS were responsible for the gene prediction and annotation. TU, WP, and CS performed repeat analysis. IN and PJ carried out comparative genome analysis. TU, SA, WR and NK performed differential expression analysis and validation. WP, IN, FT and NK supervised this project. TU, WP, IN, and NK wrote the manuscript. All authors contributed to the final manuscript editing.

