## Supporting Information for "A chromosome-level assembly of the black tiger shrimp (*Penaeus monodon*) genome facilitates the identification of novel growth-associated genes"


| | |
|---|---|
| Table S1. Published transcript sequences of *P. monodon* used for gene annotation. | Page 2 |
| Table S2. Body weight of five-month-old female shrimp in the fast-growing shrimp and the slow-growing shrimp. | Page 3 |
| Table S3. Annotation statistics for *P. monodon*. | Page 4 |
| Table S4. Differentially expressed genes between fast-growing shrimp and the slow-growing shrimp (Log2 fold-change>1 and p-value <0.05). | Excel file |
| Table S5. Differentially expressed genes mapped on KEGG pathways. | Excel file |
| Figure S1. Weight distribution of 140 sampling shrimps | Page 5 |
| Figure S2. Circular map of the mitochondrial genome of *P. monodon*. | Page 6 |
| Figure S3. COG functional categories of the differentially expressed genes in fast-growing shrimp and slow-growing shrimp. | Page 7 |
| Figure S4. Validation of RNA-seq data by quantitative real-time PCR. | Page 8 |
| Supplemental method | Page 9 |

**Table S1. Pre-processed published transcript sequences of *P. monodon* used for gene annotation.**

| Sequence Read Archive (SRA) | Paired-end reads | Total bases (bp) | Number of reads | Shortest length (bp) | Average length (bp) | Longest length (bp) |
|---|---|---|---|---|---|---|
| PRJNA4214000 | Forward | 73,136,788,946 | 599,684,773 | 100 | 122 | 125 |
| | Reverse | 73,133,977,373 | 599,684,773 | 100 | 122 | 125 |
| SRR1648423, SRR1648424 | Forward | 6,416,588,192 | 64,648,370 | 50 | 99 | 101 |
| | Reverse | 6,381,551,117 | 64,648,370 | 50 | 99 | 101 |
| SRR2191764 | Forward | 12,321,377,090 | 125,071,719 | 50 | 99 | 100 |
| | Reverse | 12,254,325,428 | 125,071,719 | 50 | 98 | 100 |
| SRR2643301, SRR2643302, SRR2643304, SRR2643305 | Forward | 11,951,482,883 | 77,980,953 | 50 | 153 | 250 |
| | Reverse | 11,891,135,923 | 77,980,953 | 50 | 152 | 250 |

**Table S2. Body weight of five-month-old female shrimp in the fast-growing shrimp and the slow-growing shrimp.**

| | No. | Body weight (g) | Hepatopancreas (g) |
|---|---|---|---|
| Fast-growing shrimp | 1 | 34.73 | 1.28 |
| | 2 | 36.45 | 1.44 |
| | 3 | 37.42 | 1.38 |
| | 4 | 32.44 | 1.29 |
| | 5 | 33.34 | 1.71 |
| | 6 | 38.6 | 1.16 |
| | 7 | 35.33 | 1.61 |
| | 8 | 36.56 | 1.59 |
| | 9 | 38.35 | 1.56 |
| | 10 | 33.77 | 1.33 |
| | 11 | 35.82 | 2.01 |
| | 12 | 38.18 | 1.93 |
| | 13 | 37.58 | 1.67 |
| | 14 | 37.68 | 1.93 |
| | 15 | 37.73 | 1.4 |
| | **Average±SD** | **36.27±1.96** | **1.55±0.26** |
| Slow-growing shrimp | 1 | 12.7 | 0.5 |
| | 2 | 13.3 | 0.4 |
| | 3 | 13.5 | 0.4 |
| | 4 | 13 | 0.6 |
| | 5 | 13.6 | 0.7 |
| | 6 | 13.8 | 0.5 |
| | 7 | 13.8 | 0.6 |
| | 8 | 14.1 | 0.6 |
| | 9 | 12.7 | 0.6 |
| | 10 | 13.2 | 0.3 |
| | 11 | 13.3 | 0.7 |
| | 12 | 13.5 | 0.5 |
| | 13 | 14.2 | 0.6 |
| | 14 | 14.34 | 0.65 |
| | 15 | 12.9 | 0.6 |
| | **Average±SD** | **13.46±0.52** | **0.55±0.11** |

**Table S3. Annotation statistics for *P. monodon*.**

| | |
|---|---|
| Number of predicted gene models | 36,538 |
| Total gene length (Mb) | 516.19 |
| Average gene size (nt) | 16,314 |
| Average number of exons/transcript | 6.8 |
| Total exon length (Mb) | 53.57 |
| Average exon length (bp) | 215 |
| GC content of exons (%) | 52.67 |
| Average number of Introns/ transcript | 5.8 |
| Total intron length (Mb) | 596.25 |
| Average intron length (bp) | 2,802 |
| GC content of introns | 37.58 |

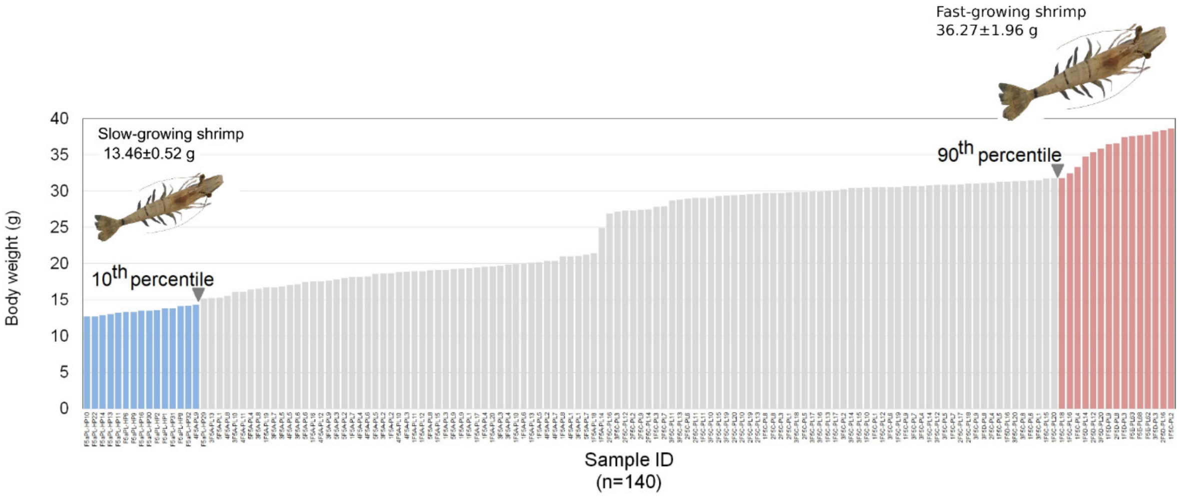


**Figure S1. Weight distribution of 140 sampling shrimps**

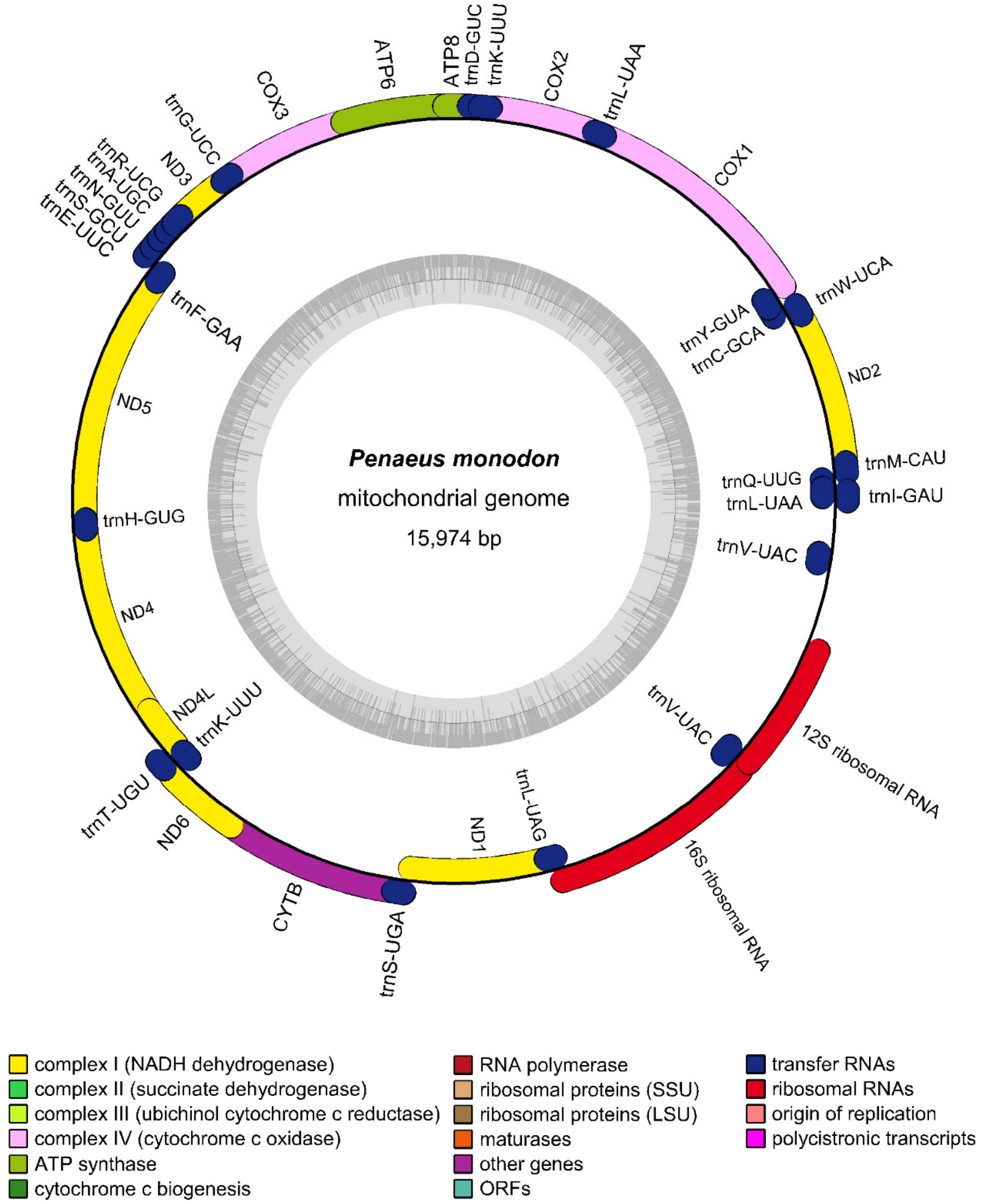


**Figure S2. Circular map of the mitochondrial genome of *P. monodon*.**

Protein coding genes, tRNA and rRNA are shown on the outer ring. Genes encoded on both strands. The inner ring shows GC density.

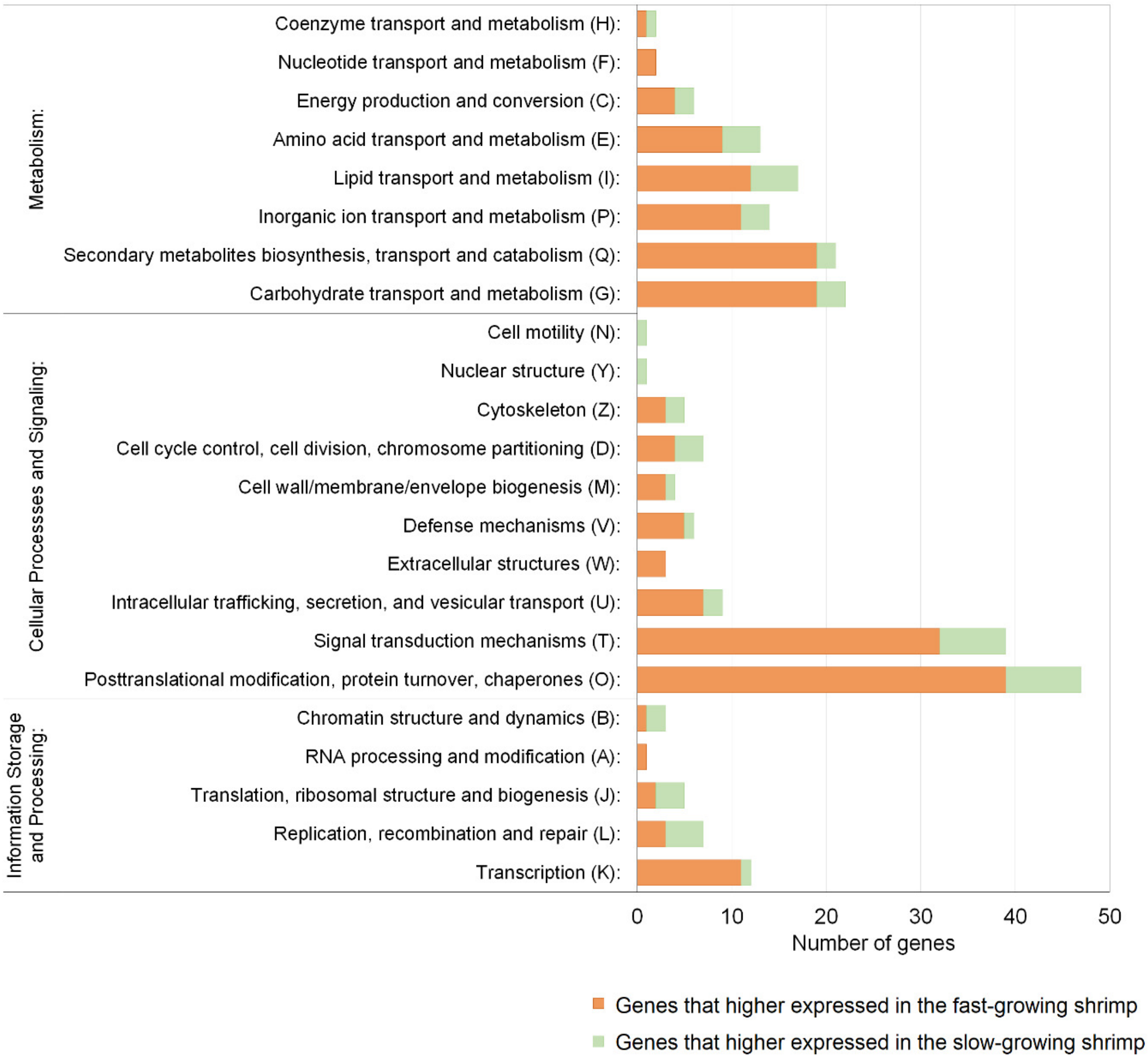


**Figure S3. COG functional categories of the differentially expressed genes in fast-growing shrimp and slow-growing shrimp**. Each bar represents the actual number of genes.

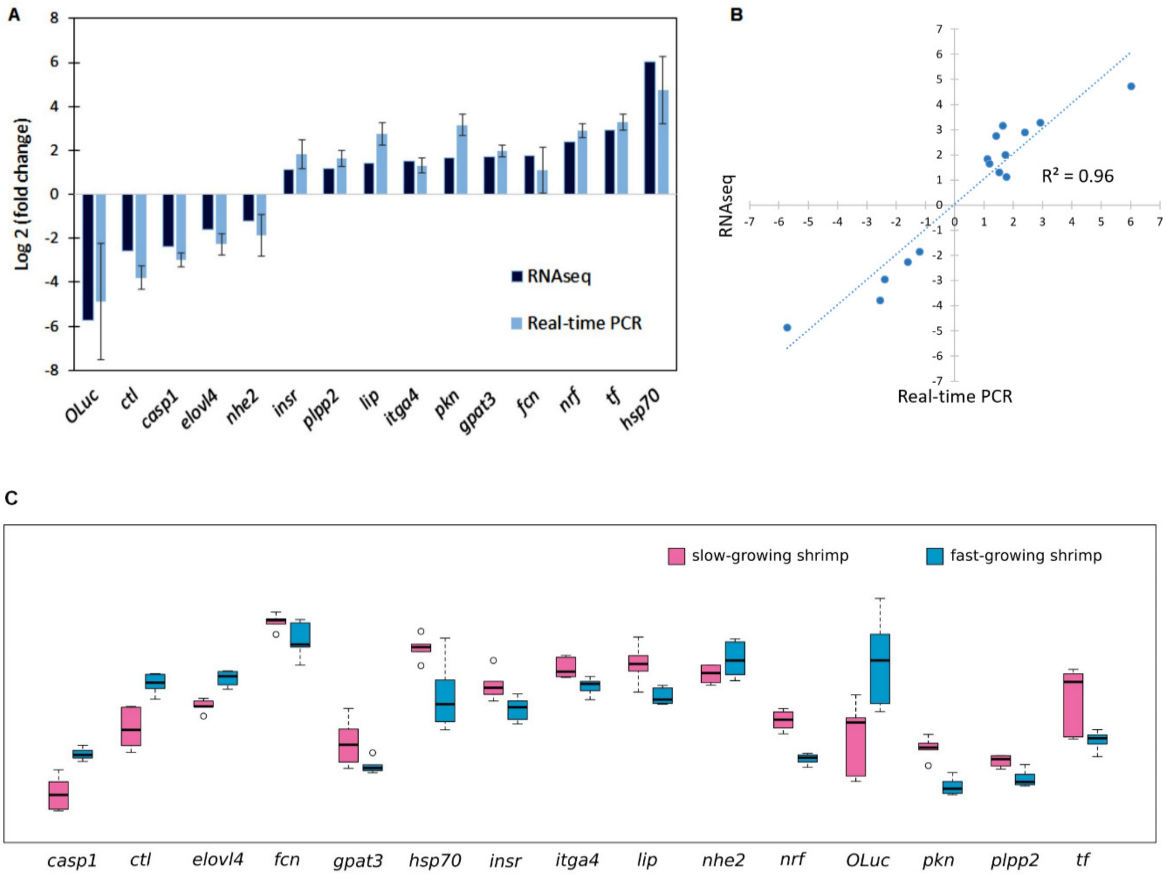


**Figure S4. Validation of RNA-seq data by quantitative real-time PCR.**

**(A)** Comparison of the expressions profile of 16 DEGs. Data shown are the mean of 8 samples ± standard deviation. **(B)** Correlation of data between RNA-seq and quantitative real-time PCR (qPCR). **(C)** Boxplot of the average expression (qPCR) from 15 genes between slow-growing and fast-growing shrimp. Validated genes included caspase-1-like (*casp1*), c-type lectin (*ctl*), elongation of very long chain fatty acids protein 4-like (*elovl4*), ficolin-1-like (*fcn*), glycerol-3-phosphate acyltransferase 3 (*gpat3*), heat shock 70 kDa protein (*hsp70*), insulin-like growth factor 1 receptor (*insr*), integrin alpha-4-like (*itga4*), lipase (*lip*), sodium/hydrogen exchanger 2 (*nhe2*), nose resistant to fluoxetine protein 6-like (*nrf*), oplophorus-luciferin 2-monooxygenase non-catalytic subunit-like (*OLuc*), serine/threonine-protein kinase N (*pkn1*), phospholipid phosphatase 2-like (*plpp2*), transferrin-like (*tf*).

### Supplemental method

#### Mitochondrial genome assembly and annotation

The PacBio reads matched mitochondria sequence (NC_002184.1) over 90% identity with the minimal 50 coverage of either the reference or the PacBio reads using BLASTN (Camacho et al. 2009) were excluded from nucleus reads and processed as following. As we observed replication of mitochondrial sequences in PacBio long reads, the reads were split into in respect to alignment match. The reads were assembled and polished using WTDBG2 (Ruan and Li 2019). Illumina reads were then aligned to the assembled contigs by minimap2 (Li 2018) for polishing using WTDBG2 using wtpoa-cns mode. The annotation was performed using the GeSeq online tool with default parameters (Tillich et al. 2017).
